# Layer 6 corticothalamic neurons induce high gamma oscillations through cortico-cortical and cortico-thalamo-cortical pathways

**DOI:** 10.1101/2024.10.05.616808

**Authors:** S. Russo, E. D. Dimwamwa, G. B. Stanley

## Abstract

Layer 6 corticothalamic (L6CT) neurons project to both cortex and thalamus, inducing multiple effects including the modulation of cortical and thalamic firing, and the emergence of high gamma oscillations in the cortical local field potential (LFP). We hypothesize that the high gamma oscillations driven by L6CT neuron activation are shaped by the dynamic engagement of intracortical and cortico-thalamo-cortical circuits. To test this, we optogenetically activated L6CT neurons in NTSR1-cre mice expressing channelrhodopsin-2 in L6CT neurons. Leveraging the vibrissal pathway in awake, head-fixed mice, we presented ramp-and-hold light at different intensities while recording neural activity in the primary somatosensory barrel cortex (S1) and the ventral posteromedial nucleus (VPm) of the thalamus using silicon probes. We found that the activation of S1 L6CT neurons induces high-frequency LFP oscillations in S1 that are modulated in frequency, but not in amplitude, across light intensities and over time. To identify which neuronal classes contribute to these oscillations, we examined the temporal evolution of firing rate in S1 and VPm. While most S1 neurons were steadily suppressed, VPm and S1 Layer 4 fast spiking (L4 FS) neurons evolved from being suppressed to facilitated within 500 ms, suggesting differential recruitment of the intracortical vs cortico-thalamo-cortical pathways. Finally, we found that LFP frequency selectively correlates with VPm firing rate. Taken together, our data suggest that L6CT neurons sculpt the frequency of S1 LFP high gamma oscillations through cortico-thalamo-cortical circuits, linking the recurrent interactions mediated by L6CT neurons to the high gamma oscillations observed across physiological and pathological conditions.

**Significance Statement:** Layer 6 corticothalamic (L6CT) neurons are strategically positioned to modulate the cortex and the thalamus allowing them to engage distinct, yet interlocked, circuits. Here we show that the activation of L6CT neurons in the mouse primary somatosensory cortex induces fast cortical oscillations through the coordinated engagement of cortico-thalamo-cortical and intracortical pathways. Our work reveals that these two L6CT-mediated pathways exert competing effects: while intracortical connections suppress cortical spiking, the activity of the cortico-thalamo-cortical loop rapidly evolves, facilitating cortical spiking. We demonstrate that the cortico-thalamo-cortical pathway operates on a faster timescale than the intracortical pathway and critically shapes cortical oscillation frequency. These findings reveal how the unique position of corticothalamic neurons allows them to flexibly and dynamically modulate the thalamocortical network.

## INTRODUCTION

Layer 6 corticothalamic (L6CT) neurons are an excitatory neuronal class with projections ramified throughout the cortex and spreading to higher and lower order thalamic nuclei. Thanks to these projections (Kirchgessner et al., 2020), L6CT neurons are strategically positioned for modulating the thalamocortical network (Bortone et al., 2014; Briggs and Usrey, 2008; Kim et al., 2014; Kirchgessner et al., 2021) and conveying top-down modulatory feedback. This feedback has been suggested to be critical for shaping signaling in the pathway, as it can enhance or suppress incoming sensory inputs and, thus, perception (Briggs and Usrey, 2009; Olsen et al., 2012). It has been proposed that this modulatory effect can be mediated by changes in the activity and synchronization at the population-level (Dimwamwa et al., 2024), manifested in local oscillations (Ray et al., 2008a) implicated in important aspects of behavior and function. Given their pivotal role in corticothalamic communication, it is particularly important to illuminate the impact of L6CT neurons on other neurons and population-level oscillations and how they contribute to electrophysiological and behavioral functions.

While multiple previous studies have investigated how L6CT neurons affect neuronal firing rates, relatively few have examined their effects at the local field potential (LFP) level. Optogenetic activation of L6CT neurons has been reported to induce cortical LFP oscillations whose frequency peaks between 60 to 250 Hz, falling in the range of activity referred to as high-gamma oscillations or the “ripples” band (Nevalainen et al., 2020; Ray et al., 2008a). Given the central role of L6CT neurons in communication between cortex and thalamus, their ability to provoke fast oscillations aligns with studies implicating high gamma oscillations in thalamocortical communication (Hu et al., 2023). Furthermore, the relationship between high gamma oscillations and thalamocortical connections provides an anatomical substrate for their critical role in attention, memory processing, coordination across areas, and cognitive tasks (Canolty et al., 2006; Gaona et al., 2011; Herrmann et al., 2004; Kucewicz et al., 2017; Ray et al., 2008b). Studies in the somatosensory pathway *in vitro* and in the auditory pathway *in vivo* (Crandall et al., 2015; Guo et al., 2017) found that the phase of these oscillations is locked to the spikes of cortical (including L6CT) and VPm neurons, thus suggesting a relationship between these oscillations and neuronal firing activity. This observation suggests that L6CT-induced oscillations may be mediated by both intracortical and cortico-thalamo-cortical circuits. Yet, the extent to which these two circuits contribute to the emergence of high-gamma oscillations is still elusive and broadens the long-standing question of whether L6CT effects are mainly mediated by their intracortical or cortico-thalamo-cortical projections (Sherman and Guillery, 1996).

Given the above, the ability of L6CT neurons to generate high gamma oscillations raises several questions: do L6CT neurons originate high gamma oscillations also in the somatosensory pathway in awake mice? How are these oscillations shaped and modulated? How do they relate to neuronal spiking? Here we hypothesize that L6CT neurons drive high gamma LFP oscillations in the primary somatosensory cortex (S1) of awake mice through the coordinated interaction of intracortical and thalamocortical pathways. We tested this hypothesis by manipulating the activity of L6CT neurons through targeted optogenetic stimulation and examining the effects on the cortical LFP, and the spiking activity of cortical and thalamic neurons. Our results show that the optogenetic activation of L6CT neurons induces high gamma oscillations in the somatosensory pathway of awake mice; while the amplitude of these oscillations is stable, their frequency is determined by the degree of L6CT activation and varies over time. To identify the neuronal circuits regulating this activity, we examined the firing rates of cortical and thalamic units, finding a similar evolution over time in the activity of VPm and layer 4 fast spiking (L4 FS) neurons that was distinctly different from other neurons in the circuit. Finally, we found that the evolution of VPm firing rate over time covaries with the frequency of S1 high gamma oscillations, thus suggesting that VPm may play a critical role in sculpting the frequency of S1 LFP oscillations.

## METHODS

### Data acquisition

In this paper we leverage the experiments used to study L6CT-induced single unit activity in Dimwamwa et al (Dimwamwa et al., 2024) to investigate how L6CT activation shapes cortical LFP oscillations and to explore how these oscillations relate to cortical and thalamic spiking activity. This dataset consists of the optogenetic stimulation of S1 in 12 NTSR1-cre (B6.FVB(Cg)-Tg(Ntsr1-cre)GN220Gsat/Mmcd, MMRRC) adult mice aged between 6 weeks and 6 months. In brief, mice were kept in a reversed light-dark cycle at 65-75°F and 40-60% humility. Mice underwent a first surgical procedure, thinning the skull above S1 (as identified by stereotactic coordinates), intrinsic imaging during whisker stimulation to map the location of the barrels associated with each whisker, were injected in the primary somatosensory cortex (S1) with 800 mL of cre-dependent AAV5-EF1a-DIO-hChR2(H134R)-eYFP.WPRE.hGH (Kirchgessner et al., 2020; Pauzin and Krieger, 2018) (UPenn Vector Core), and were habituated to the experimental rig. After at least 3 weeks from the injection, mice were anesthetized (induction: isoflurane 5%; Maintenance: isoflurane 0.5-2%) to open 2 burr holes above VPm (as identified by stereotactic coordinates) and S1 (as identified by intrinsic imaging).

The day of the experiment, mice were head-fixed in the experimental rig and the electrodes were inserted in S1 (A1×32-5 mm-25-177-A32, A1×32-Poly3-5mm-25s-177-A32, A1×64-Poly2-6mm-23s-160, or A1×64-Edge-6mm-20-177-A64, Neuronexus, Ann Arbor, MI), VPm (A1×32-Poly3-5mm-25s-177-A32, Neuronexus, Ann Arbor, MI), and TRN (Buzsaki32 or A1×32-Poly3-5mm-25s-177-A32, Neuronexus, Ann Arbor, MI). Electrophysiological data were acquired through a Cerebrus acquisition system (Blackrock neurotech, Salt Lake City, Utah) or TDT RZ2 Bioamp Processor (Tucker Davis Technologies, Alachua, FL) with a sampling frequency of either 30,000 or 24,414.0625 Hz, respectively.

An optical fiber (400 nm diameter) was connected to a blue LED source (470 nm wavelength) and positioned to shine light above the craniotomy in S1. L6CT neurons were activated using ramps-and-hold light waveforms (ramp: 250 ms; hold: 500 msec; light intensities: 8, 16, 24, and 30 mW/mm^2^). Additionally, to determine sensory responsiveness, mice underwent whisker stimulation through a galvanometer deflected with a sawtooth waveform at a velocity of 300 deg/s in the caudo-rostral direction. Whisker stimuli were administered with 10 Hz frequency in blocks of 1 second (i.e. a sequence of 10 sawtooth waveforms). Videography was acquired under infrared illumination using CCD cameras triggered at 30 or 200 Hz. The LED source, the galvanometer, and the camera were controlled through custom scripts in MATLAB and Simulink Real-Time (MathWorks, Natick, MA) at 1 kHz. For a detailed description of the experimental procedures see Dimwamwa et al. (Dimwamwa et al., 2024).

### Histology

After the final recording, mice were transcardially perfused with PBS (137 mM NaCl, 7.2 mM KCl, and 10 mM PB, VWR) and 4% PFA. The skull was removed, and the brain was extracted, post-fixed overnight in a PFA solution, and sliced to 100 μm sections using a vibratome (VT100S, Leica Biosystems Deer Park, IL). Slices were incubated with DAPI (2 mM in PBS, AppliChem, Council Bluffs, Iowa) for 15 min, mounted on slides with a 1,4-Diazabicyclo[2.2.2]octane solution (Sigma), and imaged with a confocal microscope (Laser Scanning Microscope 900, Zeiss, Germany).

### Videography processing

Videography data were analyzed using FaceMap (Syeda et al., 2022). The analysis was restricted to the region of the video with the mouse whiskers, and that region was used to compute the motion energy (Pala and Stanley, 2022; Wright et al., 2021). The time series of the motion energy was then smoothed over time and all trials with motion energy above a fixed threshold, indicating whisker movements, were excluded (Urbain et al., 2019; Wright et al., 2021), and all subsequent analyses were performed on the remaining trials (Dimwamwa et al., 2024). Thus, the data analyzed in this study were during periods of time with no appreciable self-induced whisker motion.

### Single-unit data preprocessing

All analyses were conducted in Matlab (MathWorks, Natick, MA). Recorded data were high-pass filtered (500 Hz cutoff frequency; 3^rd^ order Butterworth filter), median filtered across channels, sorted using Kilosort (Pachitariu et al., 2016), and manually curated using Phy2 (Rossant and Harris, 2013). We considered only units with signal-to-noise ratio (SNR) of the mean spike waveform >3 and less than 2% of the spikes violating a 2ms refractory period, where SNR here is defined as the peak-to-peak amplitude of the spike divided by the amplitude of the background or baseline “noise”. S1, VPm, and TRN units used for analyses were required to be sensory responsive (Dimwamwa et al., 2024; Pala and Stanley, 2022). S1 neurons were classified as regular-spiking or fast-spiking based on the mean spike waveform (trough-to-peak time greater than 0.5 ms or less than 0.4 ms, respectively) (Ganmor et al., 2010) and each neuron was assigned to a layer relative to the center of layer 4 (as identified by current source density analysis) (Sederberg et al., 2019; Sofroniew et al., n.d.). Based on the location of the center of layer 4 along the silicon probe, each channel was assigned to a layer based on the following thickness: layer 2/3: 300 μm, layer 4: 200 μm, layer 5: 300 μm, layer 6: 300 μm (Hooks et al., 2013; Pala and Stanley, 2022; Sederberg et al., 2019).

L6CT neurons were identified using the stimulus-associated spike-latency test (SALT) (Kvitsiani et al., 2013) to determine whether a particular neuron’s spiking was significantly increased by 5 ms light pulses compared to an equivalent duration pre-stimulus window. Using the 10Hz train of sawtooth whisker stimuli of 1 second, both VPm and TRN neurons were required to be significantly responsive to both the first and the last whisker stimulus of the train, as defined by a two-sided Wilcoxon signed rank test. Additionally, to ensure that the recorded neurons were not from the posteromedial nucleus, both VPm and TRN neurons were further identified by a response latency <15 ms for the first whisker stimulus and <20 ms for the last whisker stimulus (Ahissar et al., 2001; Dimwamwa et al., 2024; Liew et al., 2024). Furthermore, VPm neurons were required to have a spike waveform width >0.3 ms (Guo et al., 2017).

### Single-unit data processing

We computed peri-stimulus time histograms (PSTHs) from −750 to 750 ms around the onset of the ramp-and-hold light with bins of 1.5 ms. PSTHs were smoothed by applying a moving average filter (15 ms smoothing window, movmean function from Matlab). The mean firing rate of each neuron during the early (250-500) and late (500-750) window was computed by averaging the mean firing rate across trials in the respective window.

To perform the dimensionality reduction analysis, the average PSTH for each cell type was cut from −100 to +750 ms around the light onset and the average PSTH of the baseline was subtracted from the entire PSTH. Dimensionality reduction was performed using principal component analysis (PCA; pca function from Matlab, algorithm: ‘svd’) on the average PSTH traces. To explain 99% variance, we selected the first two components (see scree plot in Figure 3-1) and reported the weights and time course of each component.

**Figure 1.**
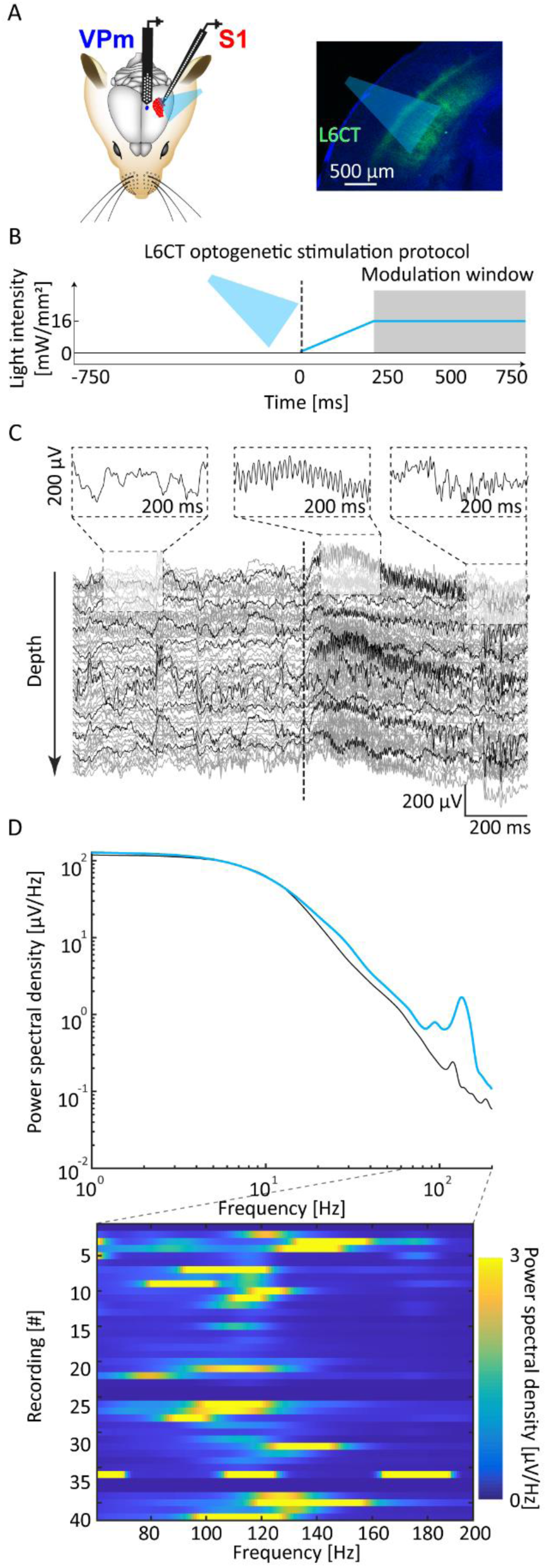
Optogenetic activation of L6CT neurons induces high gamma oscillations in the local field potential of S1. **A.** The left panel shows the schematic setup for the insertion of silicon probes in S1 and VPm during optogenetic activation of L6CT neurons. The right panel shows in blue the S1 histological section with DAPI staining and in green L6CT neurons marked with the eYFP fluorophore. **B.** The light was administered through a ramp-and-hold waveform to minimize transient effects of light. **C.** Representative traces recorded from all the silicon probe channels in S1 in one trial showing that the optogenetic activation of L6CT neurons induces high-gamma activity in the broadband local field potential (LFP). At the time of the onset of the stimulus, the LFP shows a slow deflection superimposed to high frequency oscillations. In black a subset of traces was highlighted for ease of visualization. The insets show how the oscillation changes from baseline to the beginning and end of the L6CT activation. **D.** Top: Representative broadband power spectral density (PSD) of the activity induced by the optogenetic activation of L6CT neurons during the hold-phase (light blue trace) and without optogenetic stimulation (black trace). The PSD shows that L6CT optogenetic activation generates narrowband LFP oscillations in the high gamma range. Bottom: PSD of the activity induced by L6CT activation in each recording in the 65-200 Hz band.

### LFP data processing

LFP data were downsampled to 1000 Hz (lowpass filter 200 Hz), high-pass filtered (65 Hz cutoff frequency; 3^rd^ order Butterworth filter,) and segmented from −750 to 750 ms around the onset of ramp-and-hold light manipulations.

The power spectral density (PSD) was computed using the pwelch method (pwelch function from Matlab) on the time window from 0 to 750 ms and averaged across trials. The PSD of the unfiltered signals was computed with steps of 1 Hz from 1 to 250 Hz. The PSD of the filtered signal was computed with steps of 1 Hz from 65 to 200 Hz.

To obtain the time course of the absolute voltage for each channel, we rectified the voltage and averaged it across trials. To obtain the oscillation peak frequency of each channel, we decomposed the voltage of each channel in the time-frequency domains (newtimef function from eeglab (Delorme and Makeig, 2004); Number of wavelet cycles: 5; Padratio: 8; Max frequency: 250 Hz) and identified, for each time bin and channel, the frequency showing the maximum power.

We quantified the absolute amplitude and the average peak frequency of each channel in the early and late window by averaging the absolute amplitude and the oscillation frequency, respectively, for each window. For these quantifications, we computed the Hanning windows of the wavelet filters used for the time-frequency decomposition and the moving average, and we accounted for it by reducing the early and late windows to 245-500 ms and 500-745 ms for the absolute voltage and 292-500 ms and 500-708 ms for the oscillation frequency.

### Statistical comparisons

The statistical analyses comparing the LFP amplitude and frequency across different light intensities were performed using a repeated measures ANOVA test (Muhammad, 2023). Statistical analyses comparing early and late window in units firing rate, LFP amplitude, and LFP frequency between, were performed using Wilcoxon signed-rank tests (Woolson, 2005).

The correlations between single unit PSTH and LFP amplitude and frequency were computed as follows: PSTHs were resampled to the same frequencies of LFP amplitude and frequency; signals were cut in the stable light window (250-750 ms) and correlated over time with the simultaneously recorded LFP amplitude and frequency (Spearman’s correlation) (de Winter et al., 2016).

## RESULTS

To investigate how L6CT neurons affect the LFP activity in S1, we activated L6CT neurons through targeted optogenetic manipulation while simultaneously recording extracellular electrophysiological activity across layers in S1 with silicon probes in awake mice. Specifically, we injected NTSR1-Cre mice with a recombinant adeno-associated virus (rAAV) encoding eYFP fused with Cre-dependent Channelrhodopsin2 (ChR2) in S1 (Figure 1A). To drive spiking activity in L6CT neurons, we administered blue light through a 400 μm fiber conducting light from a 470 nm LED driven with a ramp-and-hold waveform (ramp: 250ms, hold: 500ms, hold at 16 mW/mm^2^) positioned on the skull above S1 (Figure 1B). This optogenetic activation of L6CT neurons drove high frequency activity in the LFP of S1, shown across cortical depths before and during LED stimulation (Figure 1C). Qualitatively, this high frequency activity started briefly after the light onset and persisted as long as the light was on, although the oscillation pattern varied over time. Specifically, the light onset drove fast activity that progressively slowed over time. We computed the power spectral density (PSD) of the cortical LFP signal induced in the 250 to 750 ms window during L6CT activation and in periods of time with no L6CT activation of equivalent duration, finding a striking difference in the power of high frequency activity, consisting of a large narrow-band peak at approximately 150 Hz induced by the L6CT activation (Figure 1D, top). This effect was highly reproducible across recordings, but the corresponding peak frequency changed from 80 to 180 Hz depending on the subject and light intensity, as illustrated in the colormap of spectral power over this frequency band in Figure 1D (bottom).

To characterize the properties of the L6CT-induced high gamma oscillation, we high-pass filtered the signal above 65 Hz to remove line-noise and optoelectric artifacts, and separated for each time bin its amplitude, quantified as the absolute value of the voltage, from its frequency, quantified as the frequency with the maximum power in the time-frequency decomposition (Figure 2A). Inspection of the amplitude and time-frequency dynamics across all light intensities revealed that the oscillation amplitude rapidly increased, followed by a decrease over the initial light ramp (0-250 ms) and then remained stable during the hold period (250-750 ms) of the light. In contrast, the oscillation frequency progressively increased throughout the light ramp and then progressively declined during the hold period. The time-course of the oscillation amplitude and frequency averaged across subjects confirmed this pattern: the oscillation amplitude peaked early during the ramp phase and, after a brief reduction, remained stable over time; conversely, the oscillation frequency peaked at approximately 200 ms and then steadily decreased over time. Importantly, these observations confirm that the ramp phase exhibits transient effects of the stimulation that disappear during the hold phase. Interestingly, these traces suggested that distinct light intensities drove distinct LFP oscillations (Figure 2B). To investigate the difference in the oscillations induced by distinct light intensities, we averaged over time (250-750 ms) for each light intensity the oscillation amplitude and frequency and compared them across light intensities (Figure 2C). We found that while the oscillation amplitude was invariant across light intensities (Analysis of Variance – ANOVA repeated measures; p=0.07; Amplitude at 8 mW/mm^2^: 2.6 μV; Amplitude at 30 mW/mm^2^: 2.8 μV), the oscillation frequency significantly increased with higher light intensities (ANOVA repeated measures; p<0.001; Frequency at 8 mW/mm^2^: 104 Hz; Frequency at 30 mW/mm^2^: 130 Hz). Next, we examined the evolution of the oscillation over time in the hold period. To this aim, for each light intensity level and time window (early: 250-500 ms; and late: 500-750 ms, accounting for the relative Hamming windows; see methods), we averaged over time the oscillation amplitude and frequency (Figure 2D). We found that the oscillation amplitude was marginally modulated over time (Wilcoxon signed-rank test; p<0.01 in 8 mW/mm^2^ and 16 mW/mm^2^; Average amplitude at 30 mW/mm^2^: Early=2.85 μV; Late=2.82 μV), while the oscillation frequency consistently decreased from the early to the late time window (Wilcoxon signed-rank test; all p<0.001; Average frequency at 30 mW/mm^2^: Early=122 Hz; Late=136 Hz).

**Figure 2.**
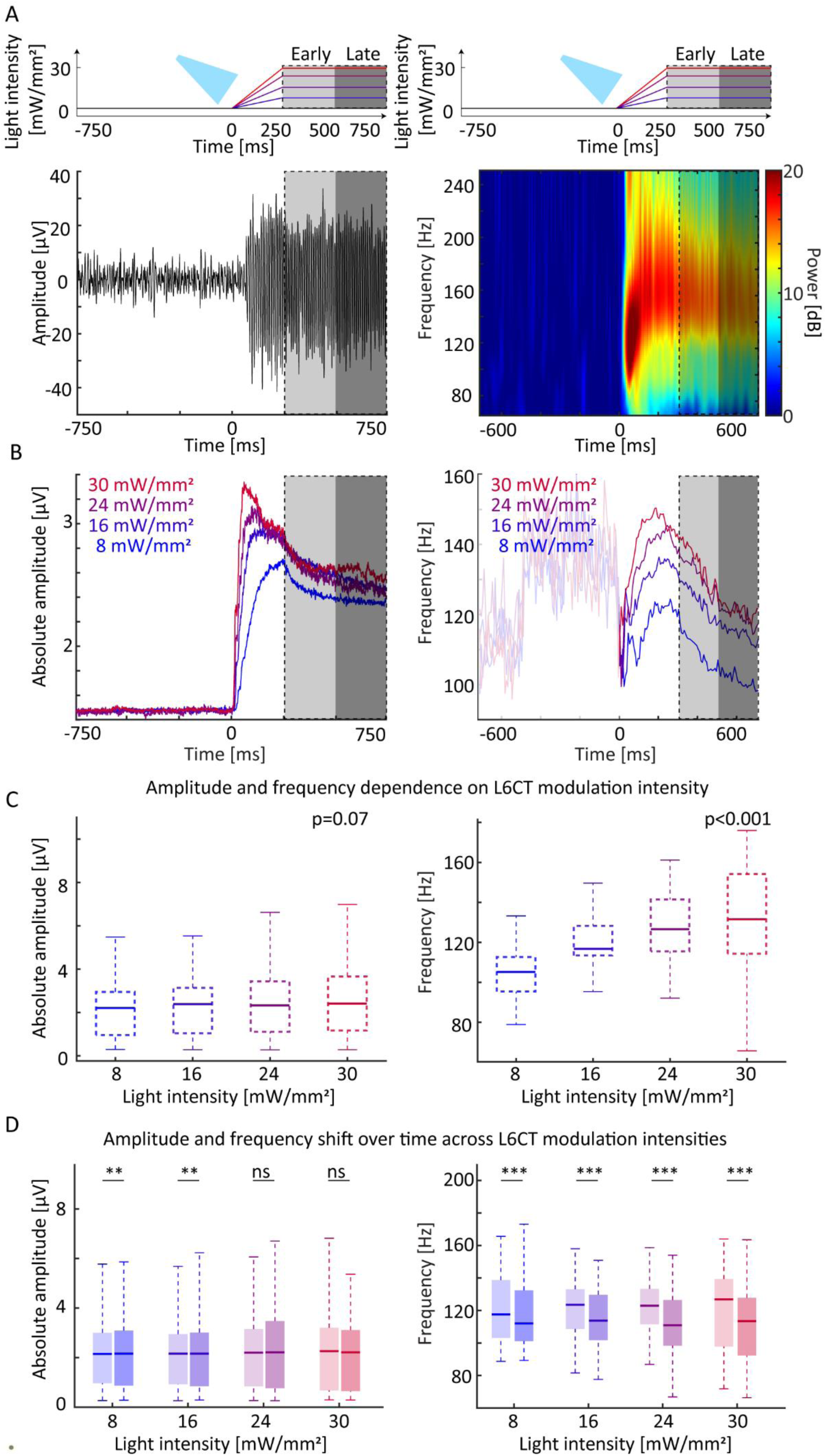
The high gamma LFP oscillations induced by L6CT neurons are intensity- and time-dependent. **A.** The top panels show that the optogenetic activation of L6CT neurons was administered through ramp-and-hold waveforms of 4 different light intensities (color coded by light intensity). The area outlined by the dashed line highlights the time window during the hold-phase of the light waveform. The shaded areas in light and dark grey highlight the early and late windows of the hold-phase of the light waveform, respectively. The bottom panels show one representative S1 LFP (left) and time-frequency decomposition (right) of the oscillation induced by the optogenetic activation of L6CT neurons showing the emergence of high gamma oscillations. The shaded regions in the bottom panel highlight smaller windows with respect to the top panels to account for the Hamming window of the filtering. **B.** The traces show the absolute voltage (left) and frequency (right; baseline window shaded) of the LFP over time across different light intensities and averaged across subjects (color code as in panel A). **C.** The boxplots show the distribution of the absolute voltage (left) and frequency (right) of the LFP during the hold-phase across different light intensities (line: median; box: 25 and 75 percentiles; whiskers: minimum and maximum value; color code as in panel A; ANOVA repeated measures). **D.** The boxplots show the distribution of the absolute amplitude (left) and frequency (right) across L6CT activation intensities between early (lighter) and late window (darker; color code as in panel A; Wilcoxon signed-rank tests).

Changes in the LFP oscillation frequency are associated with changes in the overall excitability of the circuit (Gao et al., 2017), that is known to be associated with different firing rates across distinct cell classes in the cortex (Tatti et al., 2017). Accordingly, we hypothesized that this evolution of excitability as inferred by changes in the LFP oscillation frequency would reflect in distinct modulation in the firing rates of distinct cell classes. L6CT neurons have two main projection targets: cortical neurons in deep layers (Kim et al., 2014) and thalamic neurons (including the reticular [TRN] and ventroposteromedial nucleus of the thalamus [VPm]) (Bortone et al., 2014; Sherman and Guillery, 1996). To investigate the impact of the changes in excitability reflected in the LFP on different cell classes in cortex and in sub-cortical recipients of L6CT projections, we examined the spiking activity induced by the optogenetic ramp-and-hold activation of L6CT neurons across cortical layers and electrophysiologic cell classes in the cortex (Regular Spiking [RS] and Fast spiking [FS]), VPm, and TRN. We found that cortical and VPm units were initially suppressed by L6CT activation, while TRN units were enhanced across all light intensities, as shown in our previous study in this same dataset (Dimwamwa et al., 2024) (Figure 3A). The suppression of most cortical neurons persisted throughout the L6CT activation across all light intensities. One exception to this were Layer 4 FS neurons in which, at higher light intensities, the initial suppression was followed by a remarkable and progressive increase in firing rate, to the point of becoming facilitated at the end of the light modulation with respect to their baseline. Another exception were L5 FS neurons, that were suppressed by the hold phase of the light stimulus at lower light intensities and almost not modulated at intermediate light intensities. Interestingly, VPm neurons exhibited a similar pattern to L4 FS neurons, and TRN neurons were facilitated during the light activation. To characterize this evolution of the firing rate over time, we compared the average firing rate of the early (250-500 ms) and late (500-750 ms; Figure 3B) windows of the constant light stimulation. We found a small but significant increase in the firing rates of most neuronal classes except for L6CT and L6 FS units. This increase was particularly remarkable in VPm and L4 FS neurons at higher light intensities (Increase at 8 mW/mm^2^: VPm=0.4 Hz; L4 FS=0.6 Hz; Increase at 30 mW/mm^2^: VPm=2.1 Hz; L4 FS=3.4 Hz). Contrary to most other neurons, TRN neurons were initially facilitated by L6CT activation, and this facilitation reduced over time across all light intensities (Reduction at 30 mW/mm^2^: 1.9 Hz). Interestingly, this effect was larger at higher light intensities.

**Figure 3.**
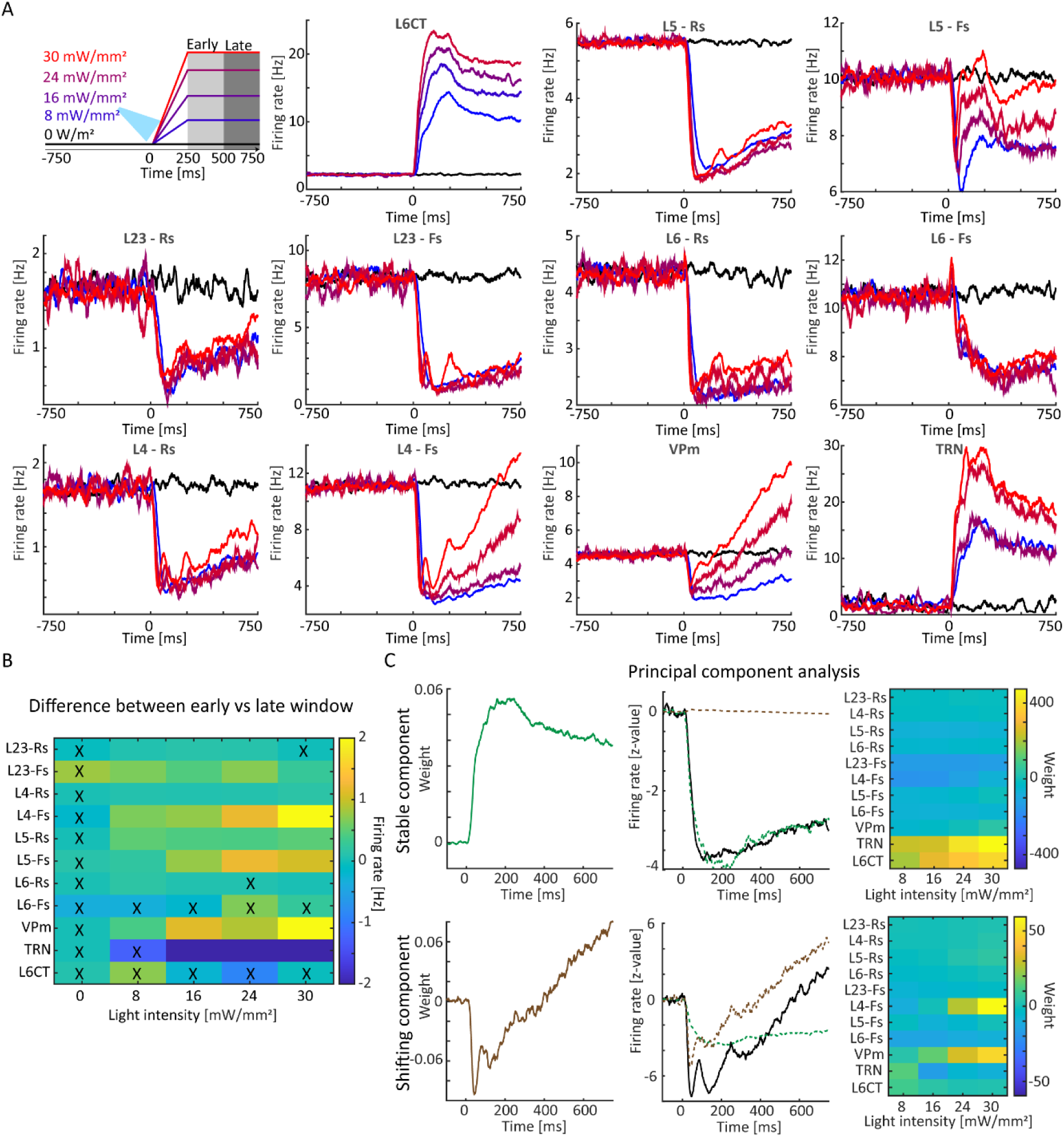
L6CT activation induces two distinct firing profiles across regions, layers, and neuronal classes. **A.** The top left panel shows the ramp-and-hold light manipulation protocol showing the time windows classified as early (light grey) and late (dark grey). The other panels show the average firing rate over time induced by ramp-and-hold optogenetic stimulation of L6CT neurons across multiple layers and classes of neurons. **B.** Comparison between the early and late firing rates across layers and neuronal classes (Wilcoxon signed-rank test; “x” indicates non-significant p>0.05). **C.** The left panels show that PCA summarizes the effect of L6CT neurons on other neurons in two components, one stable (green, top panels) and one shifting over time (brown, bottom panels), explaining ∼99% of the variance. The central panels illustrate the back projection of the components on two representative firing rate traces (L23 FS and L4 FS, respectively), showing how firing rates in different neuronal classes can either reflect only the stable component or a sum of both components. The right panels show that the stable component is represented across all cortical neuronal classes, VPm, and TRN, while the shifting component is mainly represented in VPm and L4 FS neurons.

Since L4 neurons are the main target of VPm neurons, it is possible that similarities in their firing rate patterns may reflect shared underlying dynamics. To capture these underlying dynamics, we applied Principal Component Analysis to the average time course of the firing rates of each class of neurons (PCA; Figure 3C), finding that two components were sufficient to capture ∼99% of the variance across firing rates (see scree plot in Figure 3-1). Both components were characterized by a rapid onset in the first tens of milliseconds from the beginning of the ramp, but the first component was stable over time, as indicated by the relatively constant weight throughout the early and late window, while the second component increased rapidly over time, as indicated by the inversion of the weights’ polarity from the early to the late window. These two patterns are reminiscent of the stable time course of most cortical neurons and of the rapidly shifting time course of VPm and L4 FS units, respectively, presented in Figure 3A. This was confirmed by the inspection of the weights of each component across the different cell-type classes, showing that the stable component was represented across all cortical layers, VPm and TRN, while the shifting component was selectively represented with positive weights in VPm and L4 FS units at higher light intensities. Interestingly, the stable component exhibited positive weights in L6CT and TRN neurons, and negative weights in all other neurons.

The LFP analysis indicated that L6CT activation induced an oscillation in the high gamma range whose amplitude was stable over time, while its frequency shifted during the constant light window. The analysis of single-unit firing rates indicated that most cortical units exhibited a stable modulation of their firing rates over time, while the firing rates of VPm and L4 FS units shifted from suppression to facilitation. The observation of similar LFP and single-unit dynamics, both consisting of an element stable over time and an element shifting in the timescale of 500 ms, suggested a potential relationship between them. To investigate the relation between LFP and spiking dynamics, we correlated the firing rate of each neuron with the simultaneously recorded LFP oscillation amplitude and frequency (Spearman’s correlation). For each class of neurons, we assessed the strength of the correlation by computing the percentage of neurons showing a significant correlation (p<0.05) with the LFP amplitude and the average Spearman’s Rho across neurons showing a significant correlation (Figure 4A). First, this analysis revealed that L6CT activation increases the percentage of significant correlations between the firing rate of neurons from any class and the LFP amplitude. Additionally, we found that while most neurons in each neuronal class were significantly correlated with LFP amplitude during L6CT activation, the correlation strengths were close to 0 across all light intensities, suggesting a negligible relationship between the LFP amplitude and all cell class firing rates (max: 0.12; min: −0.1; Figure 4A).

**Figure 4.**
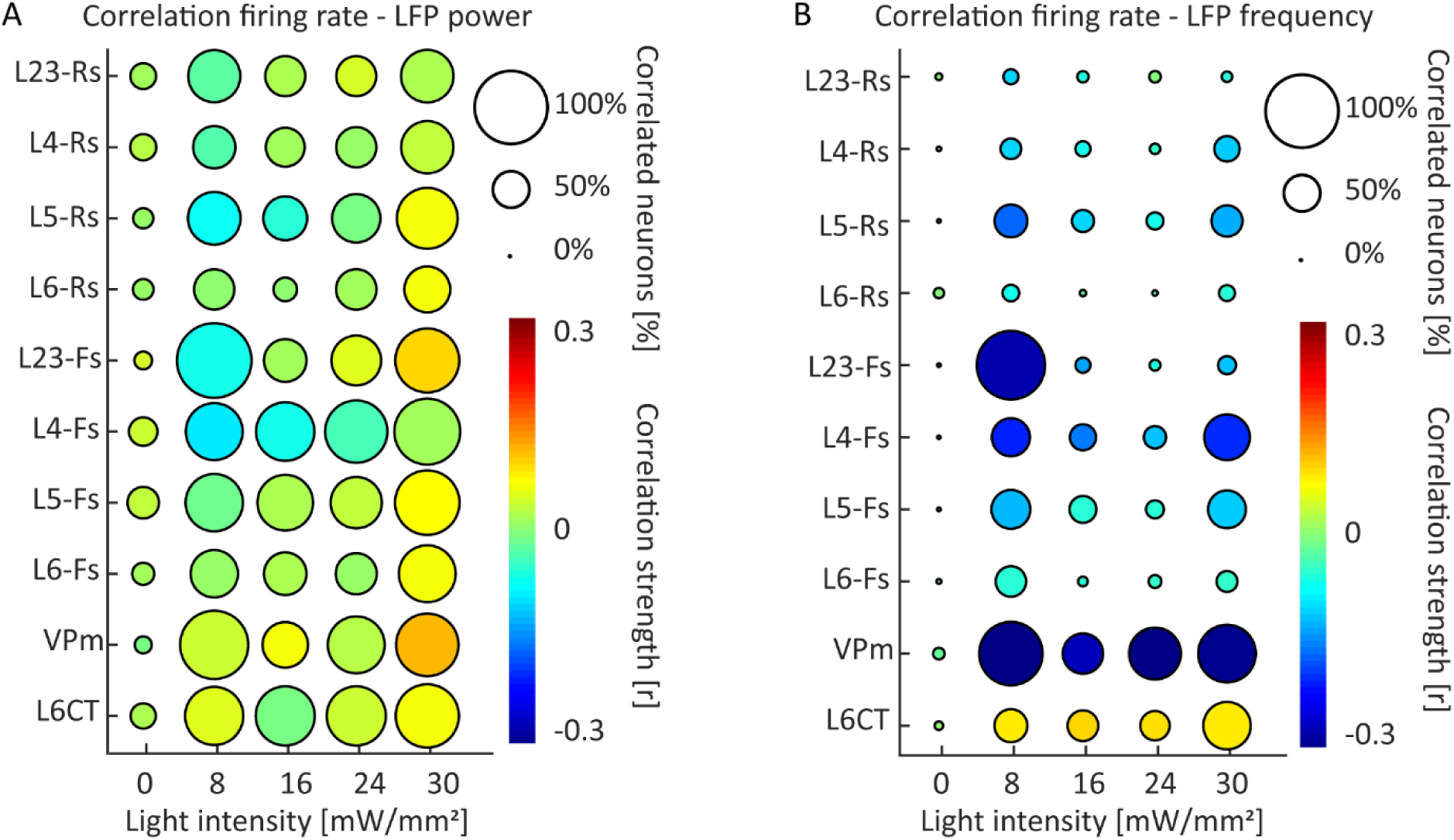
Relation between high-gamma LFP oscillation amplitude and frequency with spiking activity. **A.** The plot illustrates the relation between the firing rate of each class of neurons and the amplitude of the simultaneously recorded LFP. The circle size shows the percentage of units showing a significant correlation (Spearman correlation; p<0.05). The color code displays the average strength of the correlation, as quantified by Spearman’s Rho. The plot shows that L6CT activation induces a non-specific and weak correlation between all classes of neurons and LFP amplitude. **B.** Same as panel A for the correlation between firing rate and the frequency of the simultaneously recorded LFP. The plot shows a selective strong correlation between VPm firing rate and LFP frequency across all light intensities, and a high correlation with L23 Fs neurons at low light intensities.

We next correlated the LFP frequency with the single-unit firing rate of all cell classes in a similar manner. In contrast to the relationship with LFP amplitude, the firing rates of most cortical neurons across all classes including L4 Fs neurons were not significantly correlated with LFP frequency, except for L2/3 FS neurons at low light intensity. Another exception was the VPm neurons, as most of them were significantly and negatively correlated with LFP frequency, and with remarkable correlation strengths across all light intensities (Figure 4B). Interestingly, while a relatively small number of L6CT neurons were significantly correlated with LFP frequency, they were the only class of neurons showing a positive correlation with the LFP frequency.

## DISCUSSION

In this study, we show that L6CT neurons induce high gamma oscillations in S1 (Crandall et al., 2015; Guo et al., 2017), where the frequency of these oscillations, but not their amplitude, is a function of the level of activation of L6CT neurons and varies over time. In response to L6CT activation, the firing rate of several of the majority of excitatory and inhibitory neurons across layers exhibited an evolution in activity characterized by an initial suppression that gradually decreases, while in contrast, VPm and L4 FS neurons exhibited a remarkable switch from being suppressed to being facilitated. These spiking activity profiles were captured by two underlying components, one of which is stable over time and is represented in all neurons, and the other of which is shifting over time and is represented primarily in VPm and L4 FS neurons. By relating the spiking activity profiles of each neuronal class to LFP oscillations, we found that the firing rate of all neuronal classes was only weakly correlated with LFP amplitude, while LFP oscillation frequency was selectively and strongly correlated with VPm neuron activity.

Interestingly, we found that LFP oscillation frequency, but not amplitude, depends on L6CT activation intensity. This fine and selective modulation of the LFP frequency complements the selective modulation of LFP gamma oscillation amplitude exerted by layer 2/3 pyramidal neurons (Adesnik and Scanziani, 2010), parvalbumin neurons in the cortex (Cardin et al., 2009; Sohal et al., 2009) and in the basal forebrain (McNally et al., 2021), excitatory cortical neurons (Lu et al., 2015), hippocampal (Butler et al., 2016), and thalamocortical neurons (Hu et al., 2023). The finding that L6CT neurons can finely sculpt the frequency of high gamma oscillations suggests that they may hold a critical role in cognitive functions (Canolty et al., 2006; Gaona et al., 2011; Herrmann et al., 2004; Kucewicz et al., 2017; Ray et al., 2008b) and diseases (Mably and Colgin, 2018; McNally and McCarley, 2016; Shin et al., 2011) associated with high gamma modulations, and even more in conditions in which LFP oscillations undergo narrow-band frequency modulations, such as attention (Pulvermüller et al., 1997), seizures (Alvarado-Rojas et al., 2014; Fisher et al., 1992; Shahabi et al., 2023), dyskinesia (Wiest et al., 2022), and perception (Dubey and Ray, 2020).

One important observation here is that the temporal profile of the changes in LFP oscillation frequency correlated with the evolution of VPm firing rate. This suggests that primary thalamic nuclei may hold a key role in mediating the modulation of primary cortical LFP frequency over time. This finding aligns well with previous studies showing that the thalamus can shape the narrow-band high gamma oscillation frequency in cortical areas (Meneghetti et al., 2021) and that high gamma oscillations reflect thalamocortical communication (McAfee et al., 2018).

Our observation that at low light intensities L6CT neurons suppress downstream VPm neurons agrees with previous reports of L6CT-induced suppression of thalamic neurons *in vivo* in the visual thalamus and *in vitro* in VPm (Crandall et al., 2015; Olsen et al., 2012). This may be explained by L6CT spikes recruiting strong inhibitory TRN inputs to VPm that dominates L6CT’s direct excitatory inputs to VPm (Cruikshank et al., 2010; Dimwamwa et al., 2024). Interestingly, L6CT spikes induce a prolonged suppression of VPm at the low light intensities tested here, clearly different from the hyperpolarization that disinhibits T-type calcium channels mediating the pause-and-burst pattern observed after optogenetic VPm inhibition (Borden et al., 2022) and TRN excitation (Halassa et al., 2011; Russo et al., 2024) that has been associated with slower LFP rhythms. One potential explanation for this difference is that L6CT neurons simultaneously depolarize and hyperpolarize VPm, through their excitatory terminals or through TRN-mediated inhibition, respectively. Previous simulations suggest that the coexistence of TRN hyperpolarizing inputs and cortical depolarizing inputs may keep thalamocortical neurons in a slightly hyperpolarized state that is below spiking threshold, but above the threshold required to disinhibit T-currents and, thus, evoke bursts (Russo et al., 2024). As a result, L6CT spikes may be able to keep thalamocortical neurons in a “sweet spot” of suppression, in which reduced spiking activity can be maintained over prolonged windows of time and gradually increase, instead of abruptly switching to a bursting regime.

The finding that L6CT activation suppresses, at least at low light intensities, most cortical layers naturally raises a question about what mechanisms underlie the emergence of the cortical suppression induced by L6CT activation. Indeed, there are two mechanisms that may contribute to cortical suppression, that may coexist. The first mechanism is polysynaptic inhibition through cortical inhibitory interneurons. Previous studies have suggested that the intracortical projections of L6CT neurons in the primary visual cortex strongly recruit local inhibitory neurons which suppresses the cortex (Bortone et al., 2014; Olsen et al., 2012). Along this line, previous analyses on this same dataset indicate that L6CT activation increases the firing rate of a subset of FS neurons in S1 L4-6. An alternative mechanism of cortical suppression is the withdrawal of excitatory inputs due to suppression of VPm. Indeed, thalamic inputs are key to sustain cortical activity, and their suppression greatly reduces or even abolishes cortical firing (Russo et al., 2024; Shu et al., 2003). In support of this mechanism, our results show that one single factor captures the shift of firing activity in both VPm and L4 FS neurons, a primary target of VPm neurons. Hence, we suggest that the withdrawal of excitatory inputs from VPm contributes to the L6CT-induced suppression at least in L4 FS neurons. By contrast, the other cortical neurons mainly present stable suppression, advocating for intracortical suppression. Overall, our analyses suggest that intracortical inhibition contributes to the suppression of cortical activity across different layers throughout the entire window, while the withdrawal of excitatory VPm inputs contributes to it only for a relatively short time at the beginning of L6CT activation.

The firing rate dynamics induced by L6CT activation can be recapitulated by two principal components, with one stable component represented across all neurons and one component that shifts over time represented only in VPm and L4 FS neurons. These two different progressions over time suggest two distinct trajectories that may reflect the short-term plasticity patterns typical of intracortical and thalamic L6CT synaptic terminals, respectively. Indeed, it has been previously reported that, within 1 second, the excitatory synaptic currents induced by the activation of L6CT neurons in their intracortical terminals varied from a ∼60% increase to a ∼20% decrease, depending on the targeted layer (Kim et al., 2014). In response to similar activation inputs, L6CT terminals in VPm showed strikingly larger effects, with a ∼400% increase of fast glutamatergic currents and a drop of ∼70% in the disynaptic GABA hyperpolarizing currents (Crandall et al., 2015; Jurgens et al., 2012). While the stable component resonates with the relatively weak depression of intracortical circuits (Kim et al., 2014), the shifting component observed in VPm and L4 FS neurons resonates with the strong facilitation of excitatory synapses and suppression of inhibitory synapses described in the thalamus (Crandall et al., 2015). Indeed, our findings align with previous *in vitro* reports that L6CT activation switches, in the timescale of one second, from suppressing to facilitating VPm neurons (Crandall et al., 2015). Overall, the two factors that we isolated may recapitulate the downstream consequences of synaptic adaptation of L6CT intracortical and thalamic projections, respectively.

Interestingly, the rapidly shifting component described here is selectively expressed at the cortical level in L4 FS neurons. While the specific regain of activity in L4 neurons following the initial suppression may be explained by the strong thalamocortical projections from VPm to L4, the selective increase of firing in FS, but not RS, neurons cannot be explained by structural connectivity alone (Kim et al., 2014; Naskar et al., 2021; Sermet et al., 2019). Instead, the selective shift in firing rate of L4 FS units may be explained by their membrane and synaptic properties: PV+ neurons, the transcriptomic class associated with fast spiking activity, can spike in response to one single excitatory, fast, and robust synaptic input from VPm, while excitatory neurons require multiple synchronous excitatory inputs from VPm in order to spike (Jouhanneau et al., 2018; Sermet et al., 2019; Swadlow, 1995). As a result, the gradual increase in VPm spikes is likely to drive FS neurons more easily than RS neurons.

L6CT neurons have been implicated in many functions and behaviours (Augustinaite and Kuhn, 2020; Kirchgessner et al., 2020; Voigts et al., 2020), as they are a strategic junction between cortex and thalamus, capable of modulating both intracortical and thalamocortical loops. Furthermore, the ability of L6CT neurons to induce time- and intensity-dependent oscillations in the cortex, associated with both facilitatory and suppressive effects on downstream neurons as well as synchrony-dependent changes across cortex and thalamus (Dimwamwa et al., 2024), corroborates the idea that L6CT neurons can flexibly adapt to exert a myriad of functions. Multiple studies have suggested that L6CT neurons may be implicated in higher cognitive functions, and this hypothesis is further supported by our finding that they can sculpt the frequency of high gamma oscillations, which have been widely implicated in higher cognitive functions (Gaona et al., 2011; Kaiser and Lutzenberger, 2005; Ray et al., 2008b).

While our findings demonstrate that L6CT neurons have the capacity to modulate the firing rate of downstream units and LFP frequency, they leave an open question as to the extent to which they serve this role in natural conditions. Indeed, the role of L6CT neurons in natural conditions, such as in modulating attention levels and shaping behaviours, is still elusive. Further investigation is needed to clarify the role of L6CT neurons in their physiological range of activity. Similarly, we explored the effects of L6CT activation on the evolution of S1 LFP spontaneous activity over time, which raises the question of whether responses to sensory stimuli are differentially modulated over different L6CT activation time windows. Finally, while our study leverages correlational analyses to establish a link between VPm firing rate and LFP oscillation frequency, future experiments are needed to verify a causal link between them, to accurately disentangle the contribution of intracortical and cortico-thalamo-cortical pathways in mediating L6TC neurons effect, and to establish the role of L6CT neurons in physiologic conditions.

## REFERENCES

Adesnik, H., Scanziani, M., 2010. Lateral competition for cortical space by layer-specific horizontal circuits. Nature 464, 1155–1160. 10.1038/nature08935

Ahissar, E., Sosnik, R., Bagdasarian, K., Haidarliu, S., 2001. Temporal frequency of whisker movement. II. Laminar organization of cortical representations. J. Neurophysiol. 86, 354–367. 10.1152/jn.2001.86.1.354

Alvarado-Rojas, C., Valderrama, M., Fouad-Ahmed, A., Feldwisch-Drentrup, H., Ihle, M., Teixeira, C.A., Sales, F., Schulze-Bonhage, A., Adam, C., Dourado, A., Charpier, S., Navarro, V., Le Van Quyen, M., 2014. Slow modulations of high-frequency activity (40-140-Hz) discriminate preictal changes in human focal epilepsy. Sci. Rep. 4, 4545. 10.1038/srep04545

Augustinaite, S., Kuhn, B., 2020. Complementary Ca2+ Activity of Sensory Activated and Suppressed Layer 6 Corticothalamic Neurons Reflects Behavioral State. Curr. Biol. CB 30, 3945–3960.e5. 10.1016/j.cub.2020.07.069

Borden, P.Y., Wright, N.C., Morrissette, A.E., Jaeger, D., Haider, B., Stanley, G.B., 2022. Thalamic bursting and the role of timing and synchrony in thalamocortical signaling in the awake mouse. Neuron 110, 2836–2853.e8. 10.1016/j.neuron.2022.06.008

Bortone, D.S., Olsen, S.R., Scanziani, M., 2014. Translaminar inhibitory cells recruited by layer 6 corticothalamic neurons suppress visual cortex. Neuron 82, 474–485. 10.1016/j.neuron.2014.02.021

Briggs, F., Usrey, W.M., 2009. Parallel processing in the corticogeniculate pathway of the macaque monkey. Neuron 62, 135–146. 10.1016/j.neuron.2009.02.024

Briggs, F., Usrey, W.M., 2008. Emerging views of corticothalamic function. Curr. Opin. Neurobiol. 18, 403–407. 10.1016/j.conb.2008.09.002

Butler, J.L., Mendonça, P.R.F., Robinson, H.P.C., Paulsen, O., 2016. Intrinsic Cornu Ammonis Area 1 Theta-Nested Gamma Oscillations Induced by Optogenetic Theta Frequency Stimulation. J. Neurosci. Off. J. Soc. Neurosci. 36, 4155–4169. 10.1523/JNEUROSCI.3150-15.2016

Canolty, R.T., Edwards, E., Dalal, S.S., Soltani, M., Nagarajan, S.S., Kirsch, H.E., Berger, M.S., Barbaro, N.M., Knight, R.T., 2006. High gamma power is phase-locked to theta oscillations in human neocortex. Science 313, 1626–1628. 10.1126/science.1128115

Cardin, J.A., Carlén, M., Meletis, K., Knoblich, U., Zhang, F., Deisseroth, K., Tsai, L.-H., Moore, C.I., 2009. Driving fast-spiking cells induces gamma rhythm and controls sensory responses. Nature 459, 663–667. 10.1038/nature08002

Crandall, S.R., Cruikshank, S.J., Connors, B.W., 2015. A corticothalamic switch: controlling the thalamus with dynamic synapses. Neuron 86, 768–782. 10.1016/j.neuron.2015.03.040

Cruikshank, S.J., Urabe, H., Nurmikko, A.V., Connors, B.W., 2010. Pathway-specific feedforward circuits between thalamus and neocortex revealed by selective optical stimulation of axons. Neuron 65, 230–245. 10.1016/j.neuron.2009.12.025

de Winter, J.C.F., Gosling, S.D., Potter, J., 2016. Comparing the Pearson and Spearman correlation coefficients across distributions and sample sizes: A tutorial using simulations and empirical data. Psychol. Methods 21, 273–290. 10.1037/met0000079

Delorme, A., Makeig, S., 2004. EEGLAB: an open source toolbox for analysis of single-trial EEG dynamics including independent component analysis. J. Neurosci. Methods 134, 9–21. 10.1016/j.jneumeth.2003.10.009

Dimwamwa, E.D., Pala, A., Chundru, V., Wright, N.C., Stanley, G.B., 2024. Dynamic corticothalamic modulation of the somatosensory thalamocortical circuit during wakefulness. Nat. Commun. 15, 3529. 10.1038/s41467-024-47863-8

Dubey, A., Ray, S., 2020. Comparison of tuning properties of gamma and high-gamma power in local field potential (LFP) versus electrocorticogram (ECoG) in visual cortex. Sci. Rep. 10, 5422. 10.1038/s41598-020-61961-9

Fisher, R.S., Webber, W.R., Lesser, R.P., Arroyo, S., Uematsu, S., 1992. High-frequency EEG activity at the start of seizures. J. Clin. Neurophysiol. Off. Publ. Am. Electroencephalogr. Soc. 9, 441–448. 10.1097/00004691-199207010-00012

Ganmor, E., Katz, Y., Lampl, I., 2010. Intensity-dependent adaptation of cortical and thalamic neurons is controlled by brainstem circuits of the sensory pathway. Neuron 66, 273–286. 10.1016/j.neuron.2010.03.032

Gao, R., Peterson, E.J., Voytek, B., 2017. Inferring synaptic excitation/inhibition balance from field potentials. NeuroImage 158, 70–78. 10.1016/j.neuroimage.2017.06.078

Gaona, C.M., Sharma, M., Freudenburg, Z.V., Breshears, J.D., Bundy, D.T., Roland, J., Barbour, D.L., Schalk, G., Leuthardt, E.C., 2011. Nonuniform High-Gamma (60– 500 Hz) Power Changes Dissociate Cognitive Task and Anatomy in Human Cortex. J. Neurosci. 31, 2091–2100. 10.1523/JNEUROSCI.4722-10.2011

Guo, W., Clause, A.R., Barth-Maron, A., Polley, D.B., 2017. A Corticothalamic Circuit for Dynamic Switching between Feature Detection and Discrimination. Neuron 95, 180–194.e5. 10.1016/j.neuron.2017.05.019

Halassa, M.M., Siegle, J.H., Ritt, J.T., Ting, J.T., Feng, G., Moore, C.I., 2011. Selective optical drive of thalamic reticular nucleus generates thalamic bursts and cortical spindles. Nat. Neurosci. 14, 1118–1120. 10.1038/nn.2880

Herrmann, C.S., Munk, M.H.J., Engel, A.K., 2004. Cognitive functions of gamma-band activity: memory match and utilization. Trends Cogn. Sci. 8, 347–355. 10.1016/j.tics.2004.06.006

Hooks, B.M., Mao, T., Gutnisky, D.A., Yamawaki, N., Svoboda, K., Shepherd, G.M.G., 2013. Organization of Cortical and Thalamic Input to Pyramidal Neurons in Mouse Motor Cortex. J. Neurosci. 33, 748–760. 10.1523/JNEUROSCI.4338-12.2013

Hu, H., Hostetler, R.E., Agmon, A., 2023. Ultrafast (400 Hz) network oscillations induced in mouse barrel cortex by optogenetic activation of thalamocortical axons. eLife 12, e82412. 10.7554/eLife.82412

Jouhanneau, J.-S., Kremkow, J., Poulet, J.F.A., 2018. Single synaptic inputs drive high-precision action potentials in parvalbumin expressing GABA-ergic cortical neurons in vivo. Nat. Commun. 9, 1540. 10.1038/s41467-018-03995-2

Jurgens, C.W.D., Bell, K.A., McQuiston, A.R., Guido, W., 2012. Optogenetic stimulation of the corticothalamic pathway affects relay cells and GABAergic neurons differently in the mouse visual thalamus. PloS One 7, e45717. 10.1371/journal.pone.0045717

Kaiser, J., Lutzenberger, W., 2005. Human gamma-band activity: A window to cognitive processing. NeuroReport 16, 207.

Kim, J., Matney, C.J., Blankenship, A., Hestrin, S., Brown, S.P., 2014. Layer 6 corticothalamic neurons activate a cortical output layer, layer 5a. J. Neurosci. Off. J. Soc. Neurosci. 34, 9656–9664. 10.1523/JNEUROSCI.1325-14.2014

Kirchgessner, M.A., Franklin, A.D., Callaway, E.M., 2021. Distinct “driving” versus “modulatory” influences of different visual corticothalamic pathways. Curr. Biol. CB 31, 5121–5137.e7. 10.1016/j.cub.2021.09.025

Kirchgessner, M.A., Franklin, A.D., Callaway, E.M., 2020. Context-dependent and dynamic functional influence of corticothalamic pathways to first- and higher-order visual thalamus. Proc. Natl. Acad. Sci. U. S. A. 117, 13066–13077. 10.1073/pnas.2002080117

Kucewicz, M.T., Berry, B.M., Kremen, V., Brinkmann, B.H., Sperling, M.R., Jobst, B.C., Gross, R.E., Lega, B., Sheth, S.A., Stein, J.M., Das, S.R., Gorniak, R., Stead, S.M., Rizzuto, D.S., Kahana, M.J., Worrell, G.A., 2017. Dissecting gamma frequency activity during human memory processing. Brain J. Neurol. 140, 1337–1350. 10.1093/brain/awx043

Kvitsiani, D., Ranade, S., Hangya, B., Taniguchi, H., Huang, J.Z., Kepecs, A., 2013. Distinct behavioural and network correlates of two interneuron types in prefrontal cortex. Nature 498, 363–366. 10.1038/nature12176

Liew, Y.J., Dimwamwa, E.D., Wright, N.C., Zhang, Y., Stanley, G.B., 2024. Multiple distinct timescales of rapid sensory adapation in the thalamocortical circuit. 10.1101/2024.06.06.597761

Lu, Y., Truccolo, W., Wagner, F.B., Vargas-Irwin, C.E., Ozden, I., Zimmermann, J.B., May, T., Agha, N.S., Wang, J., Nurmikko, A.V., 2015. Optogenetically induced spatiotemporal gamma oscillations and neuronal spiking activity in primate motor cortex. J. Neurophysiol. 113, 3574–3587. 10.1152/jn.00792.2014

Mably, A.J., Colgin, L.L., 2018. Gamma oscillations in cognitive disorders. Curr. Opin. Neurobiol. 52, 182–187. 10.1016/j.conb.2018.07.009

McAfee, S.S., Liu, Y., Dhamala, M., Heck, D.H., 2018. Thalamocortical Communication in the Awake Mouse Visual System Involves Phase Synchronization and Rhythmic Spike Synchrony at High Gamma Frequencies. Front. Neurosci. 12, 837. 10.3389/fnins.2018.00837

McNally, J.M., Aguilar, D.D., Katsuki, F., Radzik, L.K., Schiffino, F.L., Uygun, D.S., McKenna, J.T., Strecker, R.E., Deisseroth, K., Spencer, K.M., Brown, R.E., 2021. Optogenetic manipulation of an ascending arousal system tunes cortical broadband gamma power and reveals functional deficits relevant to schizophrenia. Mol. Psychiatry 26, 3461–3475. 10.1038/s41380-020-0840-3

McNally, J.M., McCarley, R.W., 2016. Gamma band oscillations: a key to understanding schizophrenia symptoms and neural circuit abnormalities. Curr. Opin. Psychiatry 29, 202–210. 10.1097/YCO.0000000000000244

Meneghetti, N., Cerri, C., Tantillo, E., Vannini, E., Caleo, M., Mazzoni, A., 2021. Narrow and Broad γ Bands Process Complementary Visual Information in Mouse Primary Visual Cortex. eNeuro 8, ENEURO.0106-21.2021. 10.1523/ENEURO.0106-21.2021

Muhammad, L.N., 2023. Guidelines for repeated measures statistical analysis approaches with basic science research considerations. J. Clin. Invest. 133, e171058. 10.1172/JCI171058

Naskar, S., Qi, J., Pereira, F., Gerfen, C.R., Lee, S., 2021. Cell-type-specific recruitment of GABAergic interneurons in the primary somatosensory cortex by long-range inputs. Cell Rep. 34, 108774. 10.1016/j.celrep.2021.108774

Nevalainen, P., von Ellenrieder, N., Klimeš, P., Dubeau, F., Frauscher, B., Gotman, J., 2020. Association of fast ripples on intracranial EEG and outcomes after epilepsy surgery. Neurology 95, e2235–e2245. 10.1212/WNL.0000000000010468

Olsen, S.R., Bortone, D.S., Adesnik, H., Scanziani, M., 2012. Gain control by layer six in cortical circuits of vision. Nature 483, 47–52. 10.1038/nature10835

Pachitariu, M., Steinmetz, N., Kadir, S., Carandini, M., D, H.K., 2016. Kilosort: realtime spike-sorting for extracellular electrophysiology with hundreds of channels. 10.1101/061481

Pala, A., Stanley, G.B., 2022. Ipsilateral Stimulus Encoding in Primary and Secondary Somatosensory Cortex of Awake Mice. J. Neurosci. Off. J. Soc. Neurosci. 42, 2701–2715. 10.1523/JNEUROSCI.1417-21.2022

Pauzin, F.P., Krieger, P., 2018. A Corticothalamic Circuit for Refining Tactile Encoding. Cell Rep. 23, 1314–1325. 10.1016/j.celrep.2018.03.128

Pulvermüller, F., Birbaumer, N., Lutzenberger, W., Mohr, B., 1997. High-frequency brain activity: its possible role in attention, perception and language processing. Prog. Neurobiol. 52, 427–445. 10.1016/s0301-0082(97)00023-3

Ray, S., Crone, N.E., Niebur, E., Franaszczuk, P.J., Hsiao, S.S., 2008a. Neural correlates of high-gamma oscillations (60-200 Hz) in macaque local field potentials and their potential implications in electrocorticography. J. Neurosci. Off. J. Soc. Neurosci. 28, 11526–11536. 10.1523/JNEUROSCI.2848-08.2008

Ray, S., Niebur, E., Hsiao, S.S., Sinai, A., Crone, N.E., 2008b. High-frequency gamma activity (80–150 Hz) is increased in human cortex during selective attention. Clin. Neurophysiol. 119, 116–133. 10.1016/j.clinph.2007.09.136

Rossant, C., Harris, K.D., 2013. Hardware-accelerated interactive data visualization for neuroscience in Python. Front. Neuroinformatics 7. 10.3389/fninf.2013.00036

Russo, S., Claar, L., Marks, L., Krishnan, G., Furregoni, G., Zauli, F.M., Hassan, G., Solbiati, M., d’Orio, P., Mikulan, E., Sarasso, S., Rosanova, M., Sartori, I., Bazhenov, M., Pigorini, A., Massimini, M., Koch, C., Rembado, I., 2024. Thalamic feedback shapes brain responses evoked by cortical stimulation in mice and humans. 10.1101/2024.01.31.578243

Sederberg, A.J., Pala, A., Zheng, H.J.V., He, B.J., Stanley, G.B., 2019. State-aware detection of sensory stimuli in the cortex of the awake mouse. PLOS Comput. Biol. 15, e1006716. 10.1371/journal.pcbi.1006716

Sermet, B.S., Tru ssssssssssssssssssssssschow, P., Feyerabend, M., Mayrhofer, J.M., Oram, T.B., Yizhar, O., Staiger, J.F., Petersen, C.C., 2019. Pathway-, layer- and cell-type-specific thalamic input to mouse barrel cortex. eLife 8, e52665. 10.7554/eLife.52665

Shahabi, H., Nair, D.R., Leahy, R.M., 2023. Multilayer brain networks can identify the epileptogenic zone and seizure dynamics. eLife 12, e68531. 10.7554/eLife.68531

Sherman, S.M., Guillery, R.W., 1996. Functional organization of thalamocortical relays. J. Neurophysiol. 76, 1367–1395. 10.1152/jn.1996.76.3.1367

Shin, Y.-W., O’Donnell, B.F., Youn, S., Kwon, J.S., 2011. Gamma oscillation in schizophrenia. Psychiatry Investig. 8, 288–296. 10.4306/pi.2011.8.4.288

Shu, Y., Hasenstaub, A., McCormick, D.A., 2003. Turning on and off recurrent balanced cortical activity. Nature 423, 288–293. 10.1038/nature01616

Sofroniew, N.J., Vlasov, Y.A., Hires, S.A., Freeman, J., Svoboda, K., n.d. Neural coding in barrel cortex during whisker-guided locomotion. eLife 4, e12559. 10.7554/eLife.12559

Sohal, V.S., Zhang, F., Yizhar, O., Deisseroth, K., 2009. Parvalbumin neurons and gamma rhythms enhance cortical circuit performance. Nature 459, 698–702. 10.1038/nature07991

Swadlow, H.A., 1995. Influence of VPM afferents on putative inhibitory interneurons in S1 of the awake rabbit: evidence from cross-correlation, microstimulation, and latencies to peripheral sensory stimulation. J. Neurophysiol. 73, 1584–1599. 10.1152/jn.1995.73.4.1584

Syeda, A., Zhong, L., Tung, R., Long, W., Pachitariu, M., Stringer, C., 2022. Facemap: a framework for modeling neural activity based on orofacial tracking. 10.1101/2022.11.03.515121

Tatti, R., Haley, M.S., Swanson, O.K., Tselha, T., Maffei, A., 2017. Neurophysiology and Regulation of the Balance Between Excitation and Inhibition in Neocortical Circuits. Biol. Psychiatry 81, 821–831. 10.1016/j.biopsych.2016.09.017

Urbain, N., Fourcaud-Trocmé, N., Laheux, S., Salin, P.A., Gentet, L.J., 2019. Brain-State-Dependent Modulation of Neuronal Firing and Membrane Potential Dynamics in the Somatosensory Thalamus during Natural Sleep. Cell Rep. 26, 1443–1457.e5. 10.1016/j.celrep.2019.01.038

Voigts, J., Deister, C.A., Moore, C.I., 2020. Layer 6 ensembles can selectively regulate the behavioral impact and layer-specific representation of sensory deviants. eLife 9, e48957. 10.7554/eLife.48957

Wiest, C., Torrecillos, F., Tinkhauser, G., Pogosyan, A., Morgante, F., Pereira, E.A., Tan, H., 2022. Finely-tuned gamma oscillations: Spectral characteristics and links to dyskinesia. Exp. Neurol. 351, 113999. 10.1016/j.expneurol.2022.113999

Woolson, R.F., 2005. Wilcoxon Signed-Rank Test, in: Encyclopedia of Biostatistics. John Wiley & Sons, Ltd. 10.1002/0470011815.b2a15177

Wright, N.C., Borden, P.Y., Liew, Y.J., Bolus, M.F., Stoy, W.M., Forest, C.R., Stanley, G.B., 2021. Rapid Cortical Adaptation and the Role of Thalamic Synchrony during Wakefulness. J. Neurosci. Off. J. Soc. Neurosci. 41, 5421–5439. 10.1523/JNEUROSCI.3018-20.2021

