## Extended data, Figure 3-1 for "Layer 6 corticothalamic neurons induce high gamma oscillations through cortico-cortical and cortico-thalamo-cortical pathways"

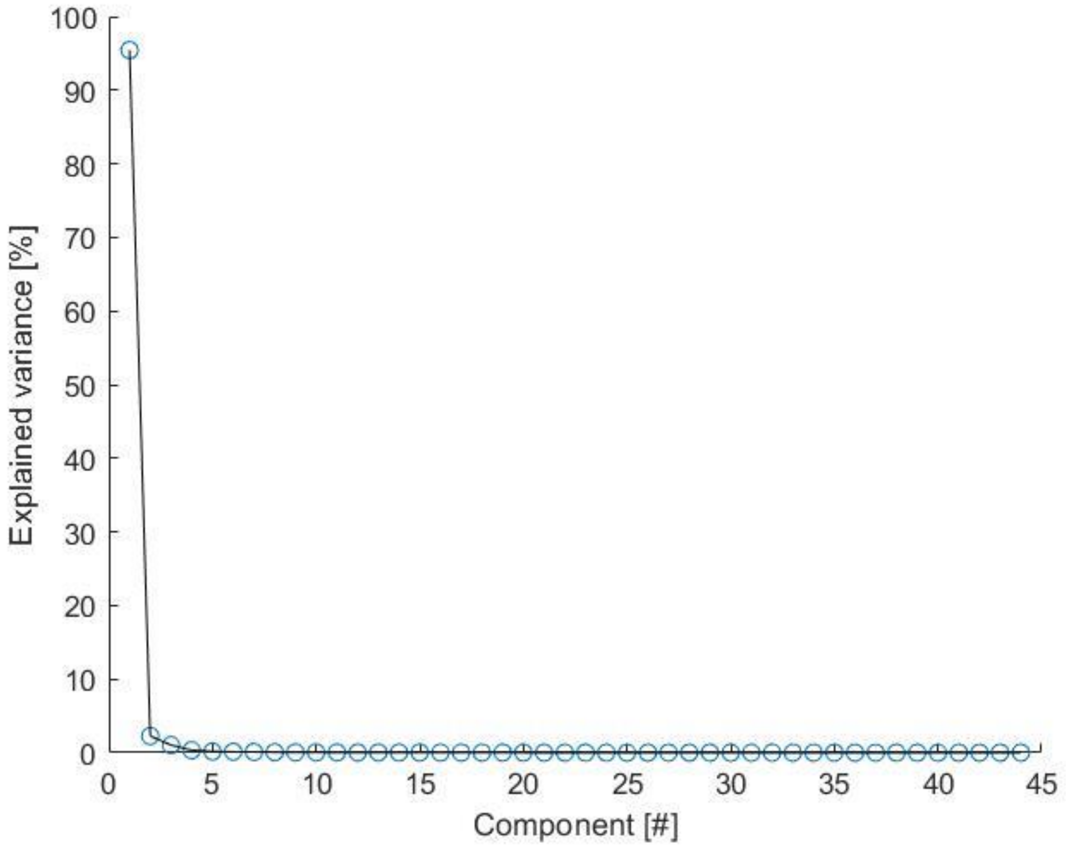


**Figure 3-1 – Scree plot for the PCA analysis.** The plot illustrates the percentual variance explained by each PCA component sorted by their explained variance.
